# Inter-brain coupling analysis reveals learning-related attention of primary school students

**DOI:** 10.1101/2022.06.08.495411

**Authors:** Jingjing Chen, Bing Xu, Dan Zhang

**Affiliations:** Dept. of Psychology, School of Social Sciences, Tsinghua University, Beijing, China; Beijing CUSoft Co., Ltd., Beijing, China; Tsinghua Laboratory of Brain and Intelligence, Tsinghua University, Beijing, China

**Keywords:** learning-related attention, EEG, primary school, inter-brain coupling, learning state monitoring

## Abstract

Learning-related attention is one of the most important factors influencing learning. While technologies have enabled the automatic detection of students’ attention levels, previous studies mainly focused on colleges or high schools, lacking further validations in primary school students. More importantly, the detected attention might fail to be learning-related if students did not attend learning tasks (e.g., the attention level of a student who reads comics secretly during classroom learning). This phenomenon poses challenges to the practical application of automatic attention detection, especially in the primary school stage, which is crucial for students to set up learning attitudes/strategies. Inspired by the emerging inter-person perspective in neuroscience, we proposed an inter-brain attention coupling method to detect learning-related attention as an extension to the existing single-person-based method. To test this method, wearable electroencephalogram (EEG) devices were used to monitor students’ attention levels in a class of primary school students during classroom learning. We found that one’s inter-brain attention coupling, defined as the degree to which an individual student’s attention dynamics match the attention dynamics averaged across classmates, was positively correlated with academic performance: higher performances are associated with higher coupling to the class-average attention dynamics. Moreover, the attention detection framework based on the inter-person perspective outperforms as an indicator of academic performance compared with the widely-used attention level within an individual. The results provide practical insights by extending the applications of detected attention levels from an inter-person perspective and demonstrating its feasibility in monitoring learning-related attention among primary school students.

## Introduction

Learning-related attention may be one of the most important factors influencing learning (Chun & Turk-Browne, 2007; Posner & Rothbart, 2014). Evidence in psychology and neuroscience suggests that attention controls what has been learned and remembered (Squire & Wixted, 2011; Weible, 2013). Nevertheless, it is difficult for teachers to spend equal time on each student to monitor their attention in the educational practice, especially in large class sizes where teachers’ time is a scarce resource (Koc & Celik, 2015; Young, 2020). For some classmates, sharing the same classroom and the same teacher does not equate to a same guidance experience (Beaman et al., 2006). Therefore, efforts have been made towards the automatic detection of students’ states in the field of education, learning science, and computer science (Dewan et al., 2019).

### EEG-based Attention Detection

Electroencephalogram (EEG) devices are able to record electrical activity in the brain from the scalp and monitor one’s attention level continuously and objectively (Davidesco et al., 2021; Sun & Yeh, 2017). Therefore, the automatic detection of students’ attention with EEG signals has received increasing interest in recent years. One line of the research focused on algorithm development to achieve high attention detection performance. For example, Hu et al. proposed an EEG-based machine learning method that combines correlation-based feature selection and a k-nearest-neighbor (KNN) algorithm. This method was able to automatically recognize three attention levels (high, low, and neutral) with an accuracy of 80.8±3.0% during a simulated distance learning task (Hu et al., 2016). Aci et al. also achieved an EEG-based attention detection (focused, unfocused, and sleep) at an accuracy of 91.72% and found support vector machine (SVM) could outperform the KNN and adaptive neuro-fuzzy system methods (Aci et al., 2019). Then, by using a convolution attention memory neural network model, Toa et al. achieved 92% accuracy, which outperformed that of the recurrent neural network, long short-term memory, and convolutional neural network. etc. (Toa et al., 2021). Al-Nafjan and Aldayel also demonstrated the feasibility of the random forest algorithm in monitoring attention levels in online learning with EEG signals (Al-Nafjan & Aldayel, 2022). These studies have shown the potentials of automatic attention detection based on EEG.

The automatically detected attention level is also expected to help teachers or online learning systems to provide individualized feedback and enhance the learning performance. For instance, Kuo et al. proposed an EEG-based attention-promoted English learning system, which is able to monitor student’s attention during English learning and triggers the attention-promoted mechanism when their attention wane by asking the student a question. Results suggested that the proposed system could increase students’ English listening achievement (Kuo et al., 2017). Chen and Wang developed an attention monitoring and alarm system based on EEG signals to help online instructors monitor the attention states of individual learners. When the system detects a low attention level for a specific student, it will alarm the student by sending an alarm message. Significantly better learning performances were observed in the group with the proposed system than in the control group (Chen & Wang, 2018). Zhang et al. proposed an EEG-based music recommendation system, which was able to recommend music based on the detected attention level to boost reading interest (Zhang et al., 2021). Gupta et al. also reported a system which used EEG signal to capture the attention levels of online leaners and provide feedback when watching online course videos (Gupta & Kumar, 2021). A recent meta-analysis further suggested that the combined effect of EEG attention detection was moderately positive in improving the online learning achievement (Liu & Zhao, 2022). Importantly, many of these researches were based on commercial EEG devices. Compared with EEG devices for research purposes, commercial EEG devices are low-cost and easy to set up. These characteristics further enhance the practical values of the above-mentioned findings and suggest an application potential in real-world educational scenarios (Lin & Kao, 2018; Xu et al., 2022). Despite these positive findings, it should be noted that most of the previous methods were explored in colleges or high schools. The feasibility of the automatic monitoring of attention levels in primary school students needs further validation. More importantly, metacognitive skills, which are crucial for lifelong learning, develop rapidly in the stage of primary school, from kindergarten age to grade six (Dignath et al., 2008). It is suggested that it is easier to change students when setting up learning attitudes/strategies than when students have already developed disadvantageous learning behaviors (Hattie et al., 1996; Hendy & Whitebread, 2000). Therefore, the automatic monitoring of attention levels to provide personalized guidance and feedback is particularly important for primary school students.

Even if it is feasible to automatically monitor attention levels among primary school students, the detected attention levels sometimes are not necessarily learning-related if students did not attend learning tasks in the first place. Imagine this: a primary school student named Bob is attending a math class in the classroom. After the bell rang, Bob secretly takes out his comics and starts reading while the teacher’s lecture is going on as background noise. Bob is not engaged in the lecture. However, we may find that the detected attention level for Bob is not lower than other students who are paying attention to the lesson. In contrast, Bob may even keep a high attention level if the comics are very interesting. Since the detected attention level is not learning-related, it will fail to characterize the learning process and even lead to wrong feedback. In previous studies, students were instructed to join in specific learning tasks (e.g., attending a lecture or reading a book) under the supervision of researchers. Under this premise, it is reasonable to assume that the detected attention level could characterize students’ learning processes during the task (Sonkusare et al., 2019). However, the educational practice in the real world is usually “wilder” and more uncontrollable compared with the experimental scenarios. The detected states may not necessarily be learning-related when students like Bob fail to attend the tasks we desired. Since primary school students have generally shown lower self-control ability than adults (Eisenberg et al., 2014), it is important to capture the learning-related attention to support teachers’ instructions and class management in primary schools.

### Inter-person Perspective in the Field of Neuroscience

The recent emerging inter-person perspective in the field of neuroscience is expected to provide a possible solution to capture the learning-related attention among primary school students (Hasson et al., 2012; Nastase et al., 2019). Unlike the present EEG-based attention detection studies that always analyze or apply attention values on a single-person level, the inter-person perspective identifies the neural correlates of interest by computing the similarity across brain patterns of a group of people who received the same stimuli or were in the same interacting scenario (e.g., the classmates or peers). Possibly the first inter-brain study reports that individual brains show a highly significant tendency to act in unison when freely viewing a popular movie. The synchronized brain activities across individuals were found in multiple brain regions, suggesting that a large extent of the human cortex is stereotypically responsive to naturalistic stimuli (Hasson et al., 2004). This finding has also been validated in typical learning scenarios such as lectures (Dikker et al., 2017), videos (Meshulam et al., 2021), and discussions (Pan et al., 2022). For example, similar brain activities are observed in students paying attention to the lesson during classroom learning (Dikker et al., 2017). Then, when students were taking a computer science course, similar brain activities were also observed in multiple brain regions across the cortex (Meshulam et al., 2021). More importantly, the shared brain responses across peers/classmates have been argued to reflect the shared attention (Dikker et al., 2017) or shared understanding (Meshulam et al., 2021) of the external stimuli (i.e., the course contents). For students situated in the same learning task, similar brain dynamics triggered by the course contents are expected to be found across students. Note that these similar brain dynamics contain not only the low-level sensory processing that closely tracks the physical features of course contents but also the higher-level functions such as semantic (Lerner et al., 2011)(Lerner et al., 2011), emotional (Hasson et al., 2004)(Hasson et al., 2004), and learning process (Meshulam et al., 2021). Then, averaging brain activities across classmates would enhance the consistent brain activities ‘tuned’ by the course contents by canceling out the idiosyncratic activities responsive to distractors. Hereby, it is plausible to consider the ‘class-average’ brain activity to represent a course-content-related learning process (see (Nastase et al., 2019) for a more detailed discussion).

Inspired by these positive findings in neuroscience, it seemed promising to consider the ‘class-average’ attention dynamics for representing the learning-related attention dynamics for primary school students. For students who are focused on the course, a high similarity is expected to be observed between their dynamics of attention levels and the class-average dynamics. For students like Bob who do not engage, we expected to observe a low similarity between his dynamics of attention levels and the class-average dynamics even if he might show a high value of attention levels. Nevertheless, no study has addressed this issue.

### Research questions

Previous studies have demonstrated the feasibility of providing individualized feedback by automatically detecting students’ attention levels in scenarios where teachers are difficult or even impossible to monitor every student’ states. However, most of the previous studies were explored in colleges or high schools. The feasibility of the automatic monitoring of attention levels in primary school students needs further validation. More importantly, the automatically detected states might fail to characterize student’s learning process if the detected state is not learning-related (e.g., the attention level of a student who reads comics secretly during classroom learning). The non-learning-related attention values may lead to useless or even wrong feedback. This phenomenon poses challenges to the practical application of the automatically detected attention in scenarios that lack the supervision of instructors, especially among primary school students who generally show lower self-control ability.

Therefore, inspired by the recent emerging inter-person approach in neuroscience, the present study aimed to propose a new attention detection framework as an extension to the existing single-person-based method. Then, we validated the framework by monitoring attention levels in a class of primary school students during their classroom learning based on a wearable EEG device. We expected to provide practical insights by extending the applications of detected attention levels from an inter-person perspective and demonstrating its feasibility in monitoring learning-related attention among primary school students.

The specific research questions were as follows:

1. How can we detect the learning-related attention level among primary school students from an inter-person perspective?
2. By taking the academic performance as the indicator, will the attention detection framework based on the inter-person perspective outperform the widely-used single-person attention metrics?

## Method

### Participants

Twenty-eight students (16 females; age: 8-9 years old) from the same class (35 students in total) from grade 3 of a primary school volunteered to wear an EEG headband during their regular class sessions. The present study was conducted in accordance with the Declaration of Helsinki. The protocol was approved by the local ethics committee. Participants, as well as their legal guardians, gave their written informed consent before joining the experiment.

#### 2.2 Procedure and Data Recording

A single-channel headband (CUBand, CUSoft, China) with dry electrodes and a NeuroSky EEG biosensor (NeuroSky, USA) was used to record EEG at Fpz (the midline of the forehead) according to the international 10-20 system, a standardized electrode placement at a sampling rate of 512 Hz (Fig.1a). The reference electrode was placed on the left ear lobe with a ground at Fp1(on the left side of the forehead). The NeuroSky EEG biosensor can calculate every second’s attention level based on their patented algorithm from EEG signals (http://neurosky.com/biosensors/eeg-sensor/algorithms/). The value for the attention level ranges from 1 to 100, indicating from “very distracted” to “very focused.” The detected attention level has been previously validated and used to monitor brain state in educational scenarios (Bitner & Le, 2022; Chen & Wang, 2018). The present study also tested the quality of the detected attention level with a meditation task as a control (see details below). Four students were omitted from the meditation data analysis as they failed to understand the meditation tasks. Fig.1b demonstrated the class-average attention dynamic in a representative session. The class-average attention level during the meditation is lower than during the lecture.

**Fig.1.**
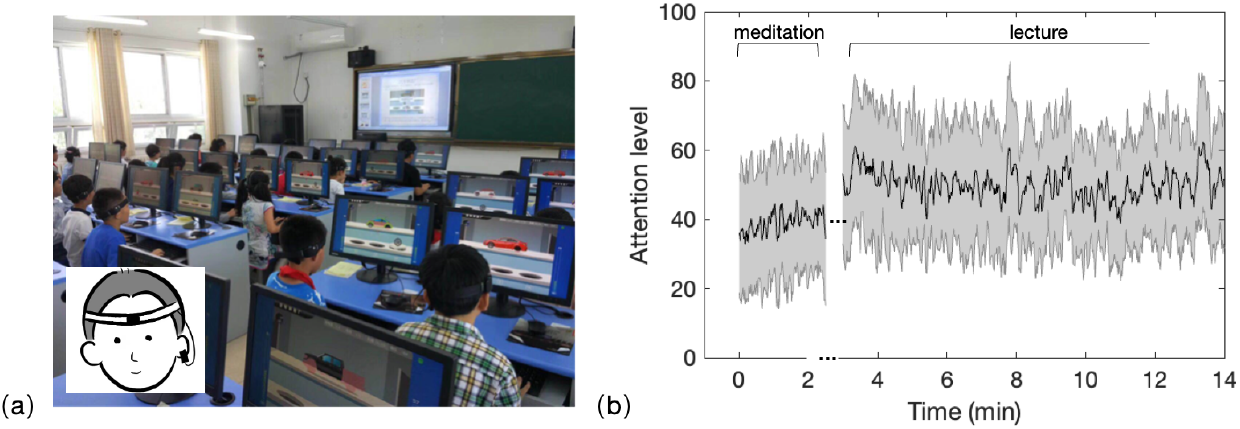
Experiment setup. (a) An illustration for students wearing EEG headbands in their classroom. (b) The time course of the class-average attention level during the meditation and lecture tasks in a representative session. The grey shadow indicated standard deviation across students at each time point.

The data collection lasted for five sessions, which were an introductory course of brain sciences designed for primary school students, covering the topics of perception, attention, emotion, memory, and language. Before each session began, students wore headbands with the help of the experimenters, and the headbands were taken off after each session. The duration of the sessions lasted around 20 minutes. During each session, students were required to meditate for around 2 minutes (close their eyes and try to relax). Then, their teacher gave a lecture.

The exam scores were used to characterize students’ academic performances. In order to exclude the possible bias in a single exam, the students’ scores in Chinese and Math on their final exam in the last term, the mid-term exam, and the final exam in the present term were averaged as the indicators of their academic performances. The mean of the students’ scores was 91.8, ranging from 74 to 99. The scores were sufficiently varied to capture the individual differences in the academic performance.

### Data Processing

To test our hypothesis, we calculated the degree of similarity between the attention dynamics of each student and the class-average attention dynamics (termed as ‘inter-brain attention coupling’) and compared its effectiveness in predicting academic performances with the widely-used attention level for each student (termed as ‘single-brain attention level’), as shown in Fig.2. For the inter-brain attention coupling, we employed an inter-subject correlation framework (Nastase et al., 2019)(Nastase et al., 2019), which has been successfully used to evaluate the learning process in a computer science lesson (Meshulam et al., 2021). First, for each student, a 30-s non-overlapping time window covering 30 attention level values was used to extract the attention dynamic during the lecture. Then, for each 30-s epoch in each student, the degree to which an individual student’s attention dynamics match the class-average attention dynamics (average across all other students) was computed using Pearson’s correlation. The correlation value was used as the inter-brain attention coupling index for each epoch. Finally, correlation values were averaged across epochs and lectures. At the same time, the single-brain attention level was obtained by averaging the attention level dynamic within and across lectures for each student. Then, there would be a single-brain index and an inter-brain index as indicators of the learning process for every student. Then, Pearson’s correlations between individuals’ inter-brain attention coupling (or single-brain attention level) and their corresponding academic performances were calculated separately. Besides, a similar analysis focused on the relaxation state was conducted as a control since the NeuroSky device could also calculate the level of relaxation in each second. The value of relaxation level ranges from 1 to 100. The detected relaxation level has also been previously applied in scenarios such as learning and stress therapy (Perhakaran et al., 2016; Ülker et al., 2017).

**Fig. 2.**
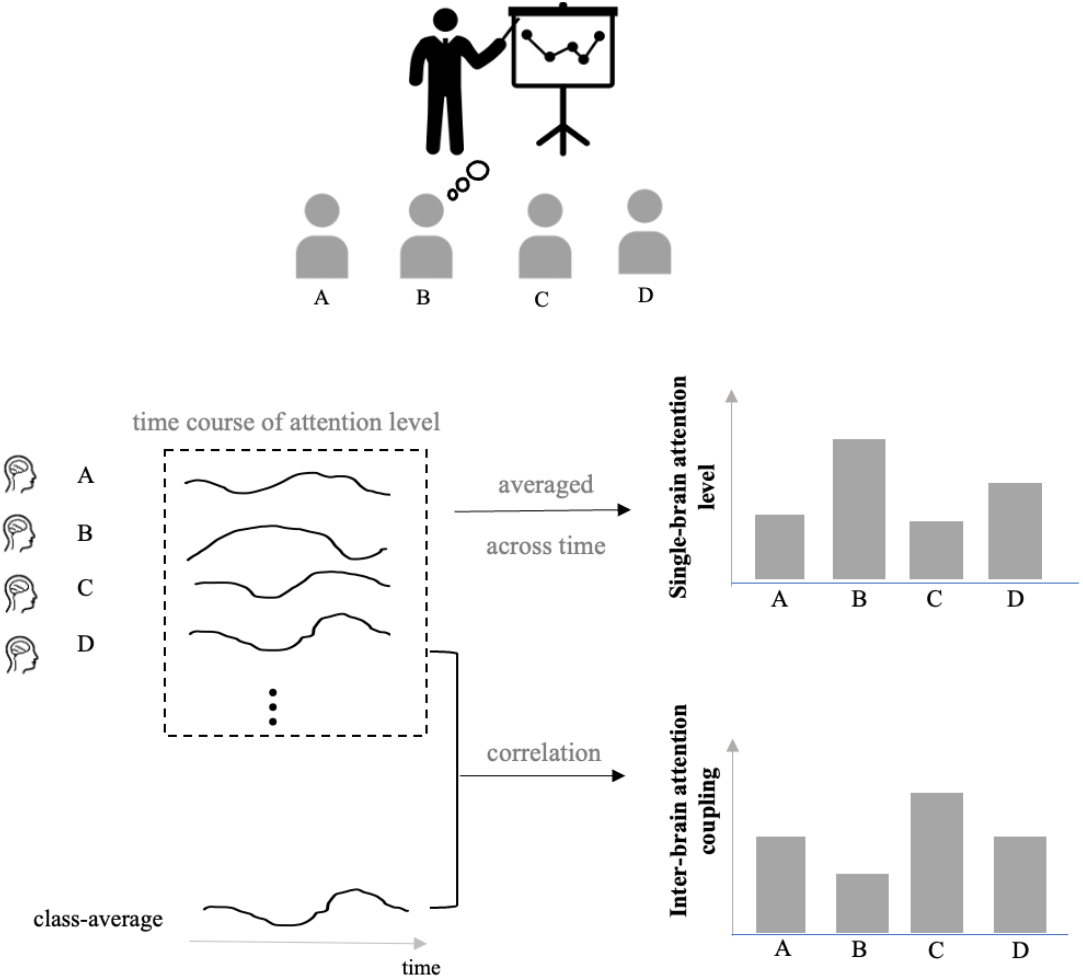
A schematic illustration of the analysis. The inter-brain attention coupling was obtained by comparing the time course of attention level for each student and the class-average pattern (average across all other students), using Pearson correlation. The single-brain attention level was obtained by averaging each student’s attention level dynamic across time.

Following the common practice of previous studies, we conducted analyses based on the EEG power in addition to the attention index. The EEG power represents the amount of brain activity in certain frequency bands of the signals (Bell & Cuevas, 2012), which has been shown to be an effective feature to characterize students’ states during learning (Al-Nafjan & Aldayel, 2022; Grammer et al., 2021; Ko et al., 2017). Therefore, the EEG power was also analyzed based on power data outputted by the NeuroSky EEG biosensor. The EEG power was calculated in each second. Here, the delta band (1-4Hz), the theta band (4-7Hz), the alpha band (8-12Hz), the beta band (13-30Hz), and the gamma band (30-40 Hz) were used in the following analysis. The power of each frequency band of interest was normalized by dividing the power in the 1-40 Hz band.

## Results

Fig.3 demonstrated the different attention dynamics during meditation and lectures. Significantly higher inter-brain attention coupling was found during the lecture task (Fig.3c, *t*(23) = 4.22, *p* = .001, paired t-test with Bonferroni correction for multiple comparison, the same below) when compared with the mediation task. We also compared the average attention level between the meditation task and the lecture. As demonstrated in Fig.3d, the single-brain attention levels were significantly higher during the lecture compared with the meditation task (*t*(23) = 5.56, *p* < .001).

**Fig.3.**
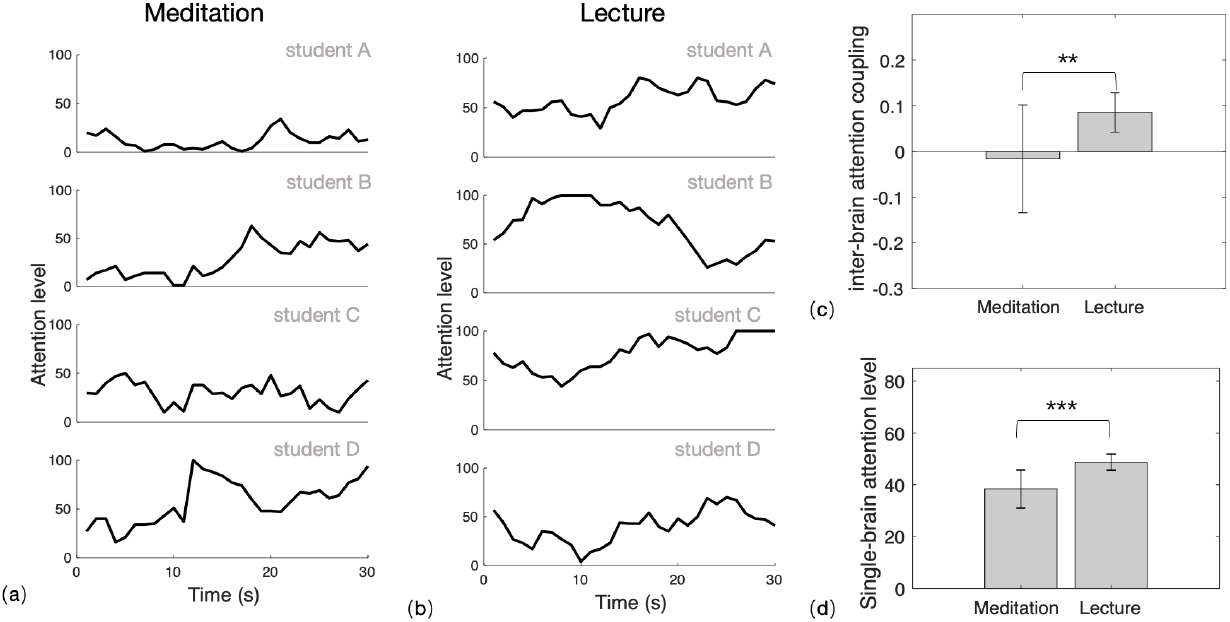
(a) The attention dynamics for representative students during meditation; (b) The attention dynamics for representative students during the lecture. (c)The comparison between the meditation task and the lecture for (c) inter-brain attention coupling and (d) single-brain attention level. *** indicated *p* < .001, ** indicated p < .01.

Here, individual’s inter-brain attention coupling during lectures was found to be positively correlated with their academic performance (Fig.4a, *r* = 0.436, *p* = .020, *n* = 28): more similar a student’s attention dynamic is to the class-average, better their academic performances are. At the same time, no significant correlations were observed between single-brain attention level and their academic performance (Fig.4b, *r* = 0.173, *p* = .379, *n* = 28). Nevertheless, the correlations in both cases increase (inter-brain attention coupling: *r* = 0.578, *p* = .003, *n* = 24; single-brain attention level: *r* = 0.384, *p* = .064, *n* = 24), respectively, when the four outliers at the left-down corner (students with academic performances < 85) were removed.

**Fig.4.**
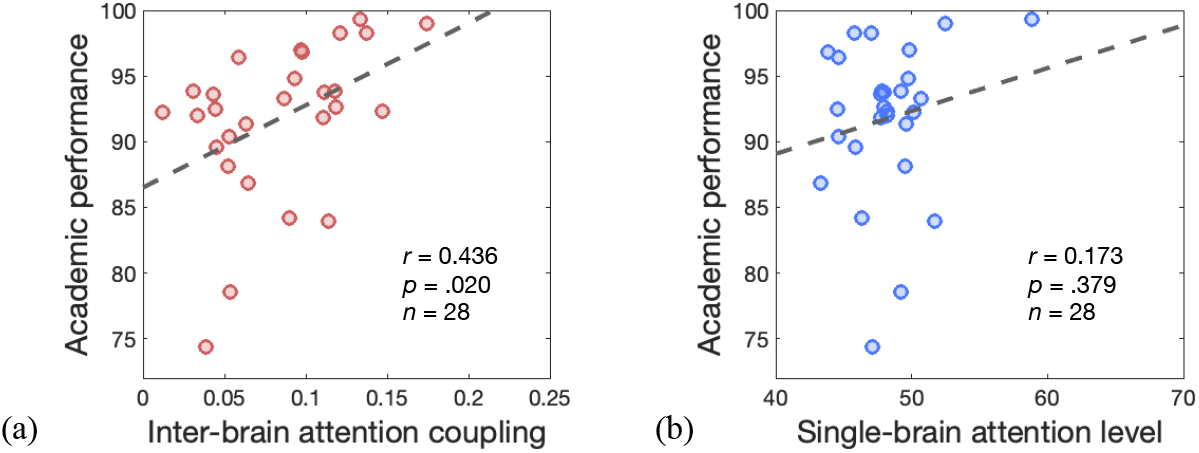
Correlations between the index for attention during lectures and academic performance. (**a**) Scatter plots between inter-brain attention coupling and academic performance. (**b**) Scatter plots between single-brain attention level and academic performance.

Fig.5 further showed the results based on the relaxation index. Single-brain relaxation levels were significantly higher in the meditation task when compared with attending the lectures (Fig. 5a, *t*(23) = 8.92, *p* < .001). As the meditation task required students to relax and decrease external attention (Rubia, 2009), the higher relaxation level (see Fig. 3d) and lower attention levels during the meditation suggested the effectiveness of these cognitive indexes. No significant difference was observed in the inter-brain relaxation couplings during the meditation task compared with the lectures (Fig. 5b, *t*(23) = 2.11, *p* > .05). Moreover, the correlations between individual’s relaxation index and academic performances also failed to reach significance (Fig. 5c-d, inter-brain relaxation level: *r* = -0.015, *p* =.940, *n* = 28; single-brain relaxation coupling: *r* = -0.278, *p* = .152, *n* = 28).

**Fig.5.**
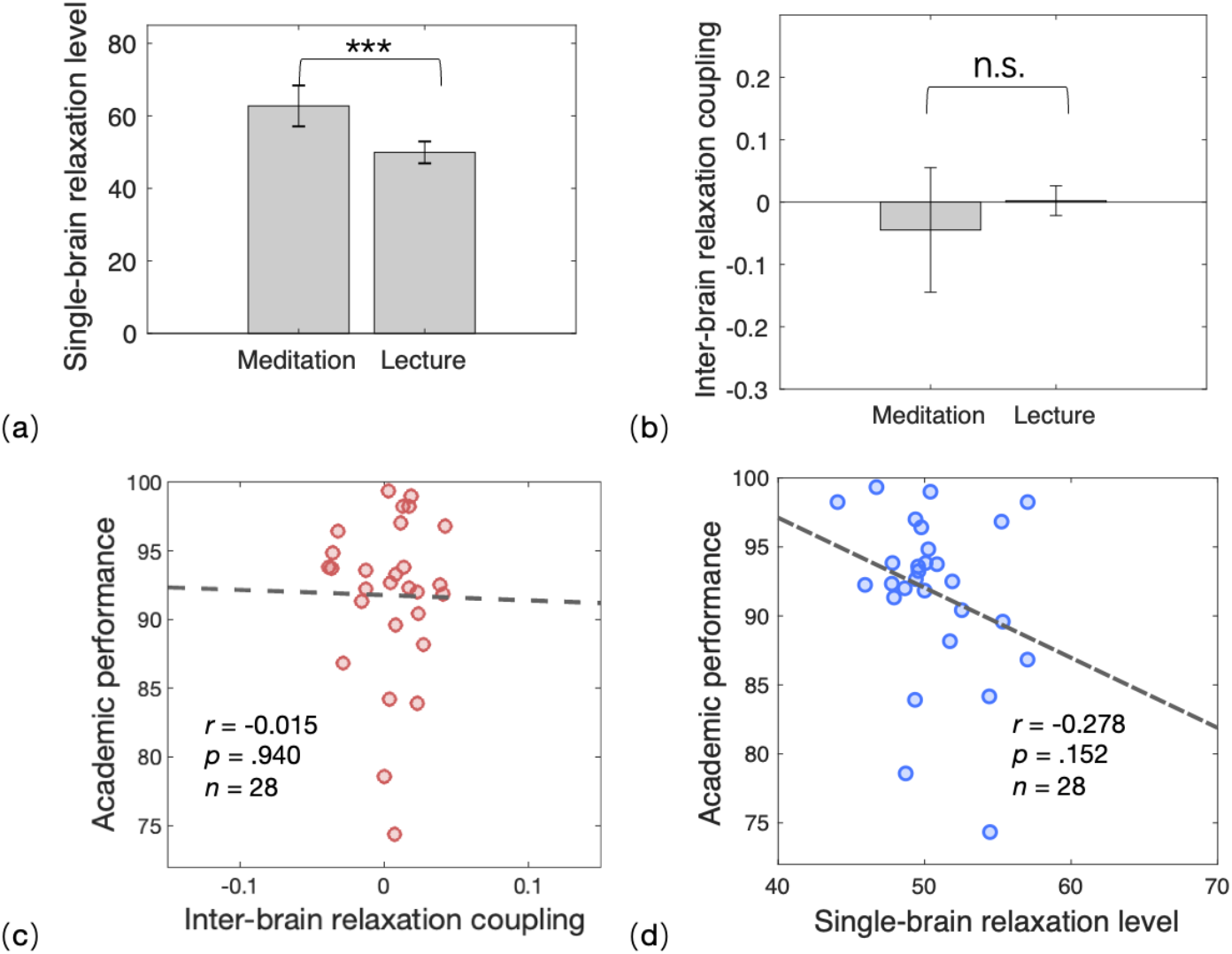
Results for the relaxation index. The comparison between the meditation task and the lecture for (a) single-brain attention level and (b) inter-brain attention coupling. *** indicated *p* < .001. n.s. indicated no significance.

Fig.6 demonstrated the summarized correlation results between students’ learning state (or EEG power) during lectures and their academic performances. As mentioned above, a significant correlation was only found between students’ inter-brain attention coupling (the similarity between a student’s attention dynamics and the average across all the other classmates) and their academic performances. When using the EEG power data for analysis, all the correlations failed to reach significance.

**Fig.6.**
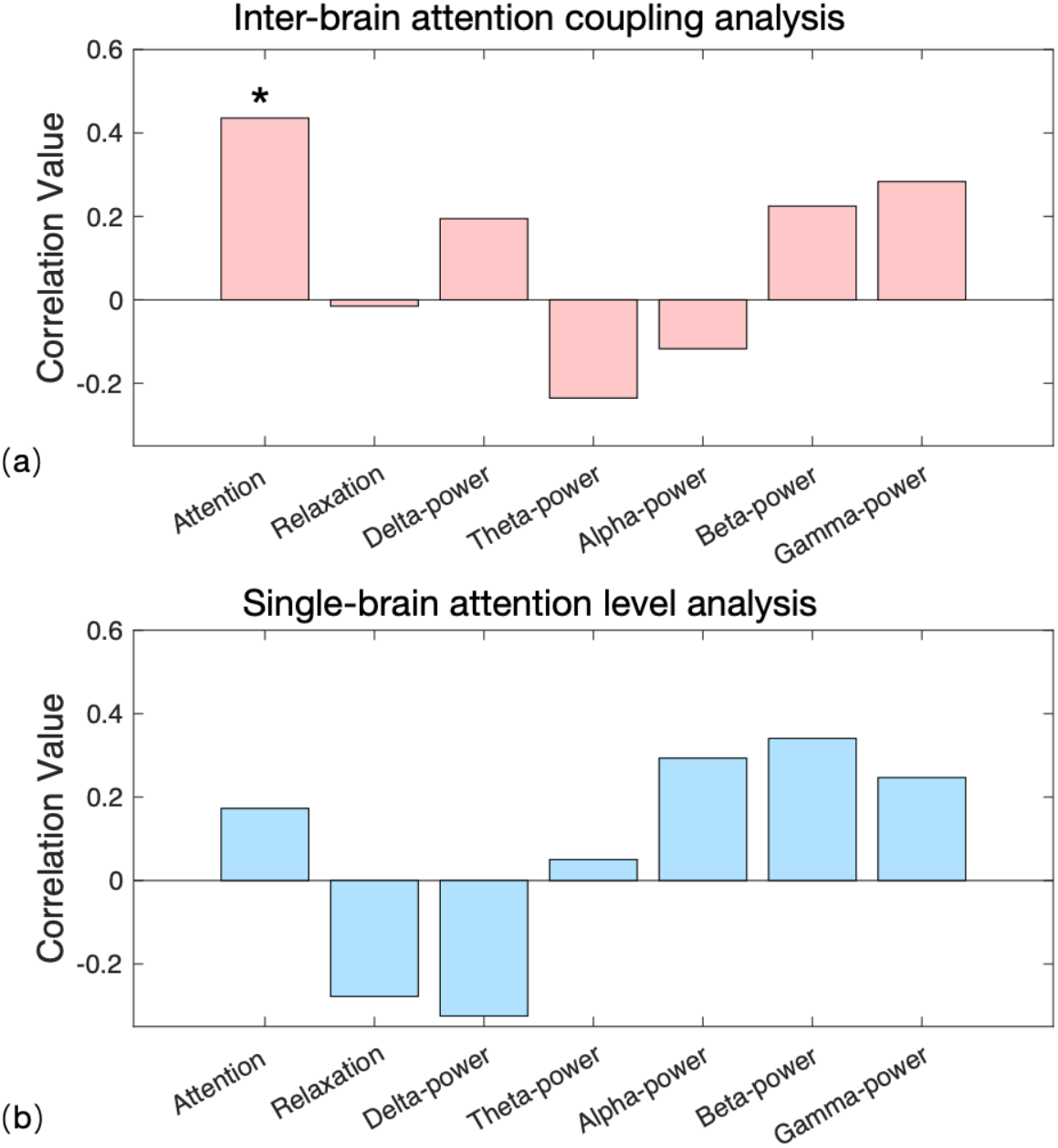
Correlation values between students’ learning state (or EEG power) during lectures and their academic performances. Bars with a pink color indicated inter-brain-based correlations, and bars with a blue color indicated single-brain-based correlations. Stars indicated a significant (*p* < .05) correlation.

## Discussion

The present study proposed an inter-brain attention coupling analysis framework to monitor students’ learning-related attention. We recorded the attention levels of a class of primary school students with wearable EEG devices during their classroom learning. Results showed that one’s inter-brain attention coupling to class-average dynamics during lectures was positively correlated with their academic performance. At the same time, no significant correlation was found between the single-brain attention level and the academic performance. A similar analysis based on the relaxation index failed to reach significant correlations. Our results verify the feasibility of the automatic monitoring of attention levels in primary school students. Then, we extend applications of detected states from an inter-person perspective and suggest its potential as a useful tool to evaluate the learning process. Compared with the widely-used attention level for individual students, we argue that inter-brain coupling analysis is expected to be particularly useful in monitoring learning states in ‘wild’ educational practical scenarios by providing learning-related information.

The present study highlighted the potential of class-average attention dynamics to represent learning-related attention dynamics during educational practices. Here, the average inter-brain attention coupling in the lecture task yielded a positive value, suggesting similar attention dynamics did emerge when students attended shared external stimuli (i.e., the course contents).This observation was further supported by the average inter-brain attention coupling in the meditation task serving as a baseline: highly individualized attention dynamics were produced when the students focused on their personalized internal states; hereby the inter-brain attention similarity was expected to be minimized, resulting in the around-zero average inter-brain coupling. Previous studies have suggested that the class-average brain dynamics could reflect the characteristics of the shared course contents (Dikker et al., 2017; Nastase et al., 2019). Then, by comparing one student’s attention dynamic with the class-average dynamic, we expected to filter out idiosyncratic dynamics responsive to distractors, thus capturing the learning-related attention dynamics intended by the course contents. Indeed, students’ academic performances were found to be significantly positively correlated with the degree of similarity between their attention dynamics and the class-average attention dynamics (i.e., inter-brain attention coupling) rather than values of the attention level (i.e., single-brain attention level), suggesting the effectiveness of the class-average attention dynamics. This finding was also consistent with previous studies where one student’s neural coupling to all their peers or classmates was positively correlated with their academic performance (Cohen et al., 2018; Meshulam et al., 2021). While previous inter-brain studies mainly focus on middle school students or undergraduate students, the present study adds evidence on the inter-brain patterns of primary school students.

The present study suggested that the inter-brain coupling analysis based on cognitive indexes calculated from brain signals could reflect the effective learning process. Compared to previous studies based on brain signals, the present results based on the attention level are expected to be more interpretable for understanding the psychological processes behind learning. Together with recent developments in cognitive state decoding based on brain signals from in the field of brain-computer interface and affective computing (Aricò et al., 2018; Gao et al., 2021), the complex mental processes during learning are expected to be further decomposed towards a deeper understanding of the real-world learning process. Using the relaxation index as a control, we highlighted the importance of learning-related attention in successful classroom learning. Besides, the present study did not observe the significant correlation between the inter-brain EEG-power coupling and academic performances, which might be explained by the different developmental stages of experiment samples, or the limited single-channel EEG recordings.

It should be noted that the non-significant single-brain-attention-level-based correlation results for the academic performances did not necessarily undermine the potential importance of the single-brain attention level. The correlation between single-brain attention level and academic performance reached marginal significance with a positive correlation coefficient of 0.384 after ruling out outliers (students with relatively low academic performances), suggesting the potential for the single-brain attention level to evaluate the learning process (Chen & Wang, 2018). Nevertheless, the correlation failed to reach significance when including all the students. One possible explanation might be that students with relatively low academic performances might focus on some distractors during class. Their single-brain attention level might not be lecture-related and eliminate the possibly-existing correlations. By contrast, the inter-brain attention coupling could be more efficient in capturing the learning-related state and correlate significantly with academic performances.

There are also limitations to be noted in the present study. First, since the present sample number is limited and focused on grade 3, further validation within a larger sample from different grades will be needed. Second, it is widely acknowledged that disciplinary differences could substantially influence learning (Neumann, 2001). More courses were needed to investigate the possible influences of disciplinary differences since there are possibly different neurocognitive mechanisms to support successful learning in different disciplines. Third, we used the attention levels and relaxation levels to describe the learning state. Although learning-related attention level has been considered the most important factor that influences the learning process (Posner & Rothbart, 2014), more diverse indexes such as mental efforts (Lin & Kao, 2018) that could be decoded from EEG are expected to provide a more comprehensive picture of students’ learning.

## Conclusion and Implications

The present study proposed an inter-person attention detection framework to monitor learning-related attention among primary school students, as an extension to the existing single-person-based method. Importantly, the present inter-person analysis framework focused on utilizing the detected attention level values, i.e., comparing values with other classmates rather than using values directly within a single person. There is no conflict between the framework with the existing EEG-based attention detection algorithms. It is possible to further enhance the detection performance for learning-related attention by combining the advanced algorithms with the proposed analysis framework.

The inter-person framework was proposed based on a classroom learning environment. Considering the validation was conducted using affordable and user-friendly commercial EEG devices, future studies are expected to facilitate a more flexible application scenario, from the classroom to online courses. First of all, the framework is readily applicable for synchronous online courses, as the inter-person analysis is not limited to the face-to-face condition and can be conducted when students are not physically present in the same location at the same time (Meshulam et al., 2021). Then, the present framework could also be expanded into the “asynchronous” condition. For example, by comparing the attention dynamics of new students with the pre-recorded attention dynamics of a group of attentive students enrolled in the same courses, it becomes possible to identify the learning-related attention levels of students who are asynchronous in online courses. Given the challenges faced by teachers in monitoring of students’ learning in online courses (Chen & Wang, 2018), further studies integrating the inter-person attention detection framework with online courses are anticipated to enhance the understanding of students’ learning states. Additionally, the high temporal resolution of EEG techniques offers the possibility for a more fine-grained description of learning-related attention dynamics. Further investigations in this direction may help teachers to identify specific course components (e.g., concept clarification or example analysis) that elicit lower levels of learning-related attention among students, thereby enabling targeted feedback and interventions.

